# Disordered breathing in a mouse model of Dravet syndrome

**DOI:** 10.1101/470609

**Authors:** Fu-Shan Kuo, Joseph L. LoTurco, Xinnian Chen, Daniel K. Mulkey

## Abstract

Dravet syndrome (DS) is a form of epilepsy with a high incidence of sudden unexpected death in epilepsy (SUDEP). Respiratory failure is a leading cause of SUDEP, and DS patients’ frequently exhibit disordered breathing. Despite this, mechanisms underlying respiratory dysfunction in DS are entirely unknown. We found that mice expressing a recurrent *SCN1a* missense mutation (A1783V) conditionally in inhibitory neurons (*SCN1a*^A1783V/+^; mixed C57B/6 background) exhibit spontaneous seizures, die prematurely and present a respiratory phenotype similar to DS patients including hypoventilation, apnea and a diminished ventilatory response to CO_2_. At the cellular level in the retrotrapezoid nucleus (RTN), we found inhibitory neurons expressing the *SCN1a* A1783V variant are less excitable, whereas chemosensitive RTN neurons, which are a key source of the CO_2_/H^+^-dependent drive to breathe, are hyper-excitable in slices from SCN1a^A1783V/+^ mice. These results show loss of *SCN1a* function can disrupt brainstem respiratory control including at the level of the RTN.

---

Dravet syndrome DS (aka. severe myoclonic epilepsy of infancy) is a severe form of early-onset epilepsy that is resistant to anti-epileptic drugs and has a high incidence of sudden unexpected death in epilepsy (SUDEP) (Kalume 2013; Kearney 2013; Shmuely *et al*. 2016). The cause of death in DS patients is thought to involve seizure-induced parasympathetic suppression of cardiac activity (Kearney 2013; Kalume *et al*. 2013; Gataullina and Dulac 2017). However, recent evidence suggests that respiratory dysfunction contributes to SUDEP associated with DS, as patients exhibit breathing problems including hypoventilation and apnea prior to the manifestation of bradycardia, a slower than normal heart rate (Kim *et al*., 2018). Patients with DS also exhibit a blunted ventilatory response to CO_2_ (Kim *et al*., 2018). This finding suggests that respiratory dysfunction, possibly at the level of respiratory chemoreceptors (neurons that regulate breathing in response to changes in tissue CO_2_/H^+^), contributes to the pathology of DS. Despite this, mechanisms underlying respiratory dysfunction in DS or epilepsy in general are entirely unknown. Leading hypothesis propose that seizure activity disrupts respiratory control by a feed-forward mechanisms involving spreading depolarization (Aiba and Noebels 2015) or activation of inhibitory subcortical projections to brainstem respiratory centers (Dlouhy *et. al*., 2015; Lacuey *et al*., 2017); however, a yet unexplored additional possibility is that epilepsy-associated mutations may directly affect brainstem respiratory centers and serve as a common substrate for both seizure and respiratory dysfunction.

Most DS cases (70-95%) are caused by mutations in the *SCN1a* gene (MIM#182389), which encodes the pore-forming subunit of a voltage-gated Na^+^ channel (Nav1.1) (Meisler and Kearney 2005; Fujiwara 2006; Catterall et al. 2010; Akiyama et al. 2012). Approximately 700 different *SCN1a* mutations have been identified in DS patients, the majority of which are missense or frameshift mutations that result in loss of function (Parihar and Ganesh 2013). Consistent with this, conventional *SCN1a* knockout mouse models (in a mixed C57B/6 background) recapitulate characteristic features of DS, including motor problems, seizures and premature death, in a remarkably titratable manner. For example, homozygous SCN1a knockout mice develop ataxia and die at 15 days postnatal, whereas heterozygous SCN1a deficient mice show seizure activity and early mortality starting at 3 weeks of age (Yu et. al., 2006; Ogiwara et al., 2007). The cellular basis for many features of DS including seizures and premature death appears to involve disinhibition, as global deletion of *SCN1a* suppresses activity of inhibitory but not excitatory neurons in the cortex and hippocampus (Yu et al. 2006; Dutton et al. 2013), and conditional deletion of *SCN1a* from forebrain inhibitory neurons results in a DS-like phenotype similar to global *SCN1a* deletion (Cheah et al. 2012). For these reasons, most studies have used global or inhibitory neuron-specific *SCN1a* deletions to model DS (Catterall WA. 2012), with few studies focusing on other high-priority genetic risk factors like *SCN1a* missense mutations, which represent ~40% of DS-associated mutations (Depienne et al., 2009; Parihar and Ganesh 2013). Thus, the extent to which expression of *SCN1a* loss-of-function mutations recapitulate features of DS remains unclear. Furthermore, despite the lethality associated with SCN1a mutations, nothing is known regarding how loss of *SCN1a* affects brainstem respiratory centers.

The main goal of this study was to provide the first detailed characterization of breathing in a SCN1a missense mutation mouse model of DS. We modeled DS by expressing a loss-of-function missense mutation (A1783V) conditionally in inhibitory neurons (referred to as *SCN1a*^A1783V/+^ mice). The A1783V variant is a recurrent DS mutation (Marini et al. 2007, Lossin 2009, Klassen et al. 2014) predicted to result in loss of function by increasing Nav1.1 voltage-dependent inactivation. Consistent with other DS models (Yu et al. 2006; Kalume et al. 2013; Kim et al. 2018), we found that *SCN1a*^A1783V/+^ mice exhibited spontaneous seizure activity and premature death starting at ~2 weeks of age, confirming this is a model of SUDEP in DS. At this same developmental time point, *SCN1a*^A1783V/+^ mice hypoventilated, exhibited frequent apneas under baseline conditions and showed a reduced ventilatory response to CO_2_, thus recapitulating the respiratory phenotype exhibited by DS patients (Kim et al. 2018). At the cellular level in a key brainstem respiratory chemoreceptor region known as the retrotrapezoid nucleus (RTN), we found that inhibitory neurons expressing A1783V show less spontaneous activity and a diminished ability to maintain firing during sustained depolarization. This is consistent with the possibility that A1783V increases Nav1.1 channel inactivation. Also consistent with a brainstem disinhibition mechanism, we found that basal activity and CO_2_/H^+^-sensitivity of excitatory chemosensitive RTN neurons was enhanced in slices from *SCN1a*^A1783V/+^ mice. These results show that RTN chemoreceptor function is altered in this DS model and may contribute to premature death.

## RESULTS

### SCN1a^A1783V/+^ mice have spontaneous seizures and die prematurely

We first sought to determine whether inhibitory neurons from SCN1a^A1783V/+^ mice express SCN1a channel transcript. We prepared brainstem sections containing the RTN from control and SCN1a A1783V/+ mice (15 days old) for subsequent fluorescent *in situ* hybridization using probes for 1) *SCN1a*, which does not distinguish SCN1a channel variants; 2) vesicular GABA transporter (VGAT) to identify GABAergic and glycinergic inhibitory neurons; and 3) vesicular glutamate transporter 2 (VGLUT2) to identify excitatory glutamatergic neurons, including chemo-sensitive RTN neurons. We labeled all cell nuclei with DAPI. Inhibitory VGAT^+^ cells were present in the RTN region and were in close proximity to excitatory VGLUT2^+^ neurons (i.e., putative RTN chemoreceptors). Both genotypes showed similar relative distributions of VGAT+ cells (T_172_ = 0.142, p = 0.88). We also observed numerous bright fluorescent puncta, which corresponded to SCN1a transcript in the soma of VGAT+ cells and, to a lesser extent, in VGLUT2+ cells in slices from control mice (F3,321 = 24.07, p < 0.0001). In slices from SCN1a^A1783V/+^ mice, we found a modest reduction in SCN1a transcript in VGAT+ but not VGLUT2+ cells (F3,321 = 24.07, p < 0.05; see Figures 1E-F). This result suggests this mutation may compromise channel expression. In a separate experiment to validate cell-type-specific Cre expression, we crossed VGATcre mice with a Rosa26^TdTomato^ reporter line and found that all tdT-labeled cells expressed VGAT, but not VGLUT2, mRNA (not shown). These results suggest inhibitory neurons from SCN1a^A1783V/+^ mice express SCN1a transcript, albeit at modestly reduced levels. Therefore, given that heterozygous deletion mutations can give rise to severe forms of DS (Yu et al. 2006, Miller et al. 2014), we expected SCN1a^A1783V/+^ mice to exhibit a mild epilepsy-like phenotype. Contrary to this expectation, SCN1a^A1783V/+^ mice exhibited a severe SUDEP-like phenotype. SCN1a^A1783V/+^ mice were born in the expected ratios, were viable, and by ~15 days postnatal, were similar in terms of body weight (T_46_ = 1.62, p = 0.11) and temperature (T_26_ =0.77, p = 0.44) as their SCN1a^+/+^ littermates (Figures 1A-C). However, SCN1a^A1783V/+^ pups showed seizure-like behavior by ~2 weeks of age (Table 1). More specifically, based on the Racine seizure-behavior scoring paradigm, only 22.7% of control mice (N = 22) showed seizure-like behavior, which mainly manifested as head-bobbing (category 1). By contrast, 77.3% of SCN1a^A1783V^ mice (N = 22) showed severe seizure behavior, including forelimb tremor (category 3), rearing alone (category 4) or in conjunction with falling over, and full-body tonic-clonic seizure (category 5). Unlike control animals, several of the mutant mice exhibited behavioral arrest that we consider to be absence seizure-like activity. In conjunction with seizure-like behavior, SCN1a^A1783V^ mice also started dying unexpectedly, reaching 100% lethality by 23 days postnatal (Figure 1D).

**Figure 1.**
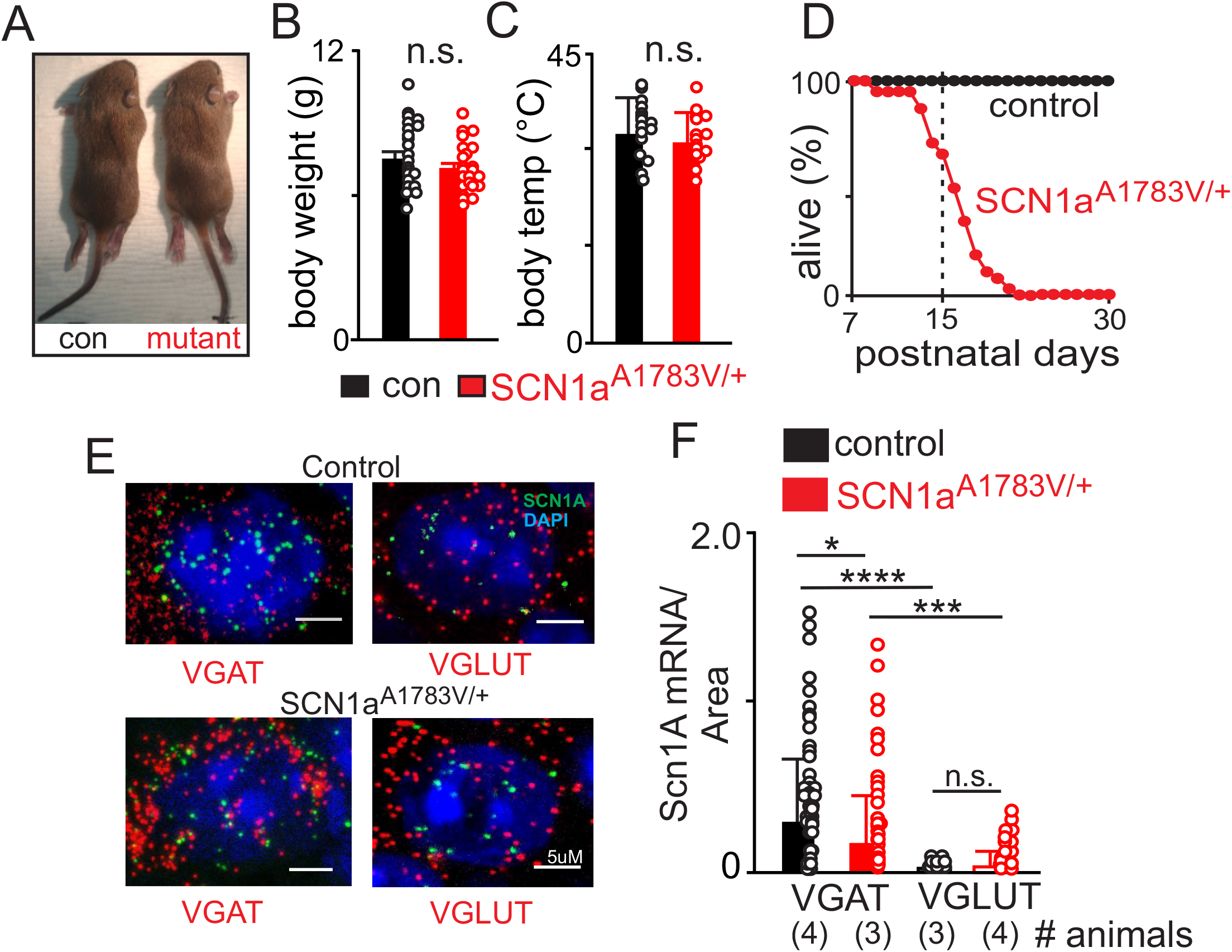
Conditional expression of SCN1a^A1783V/+^ in inhibitory neurons results in premature death. **A-C**, SCN1aA1783V/+ mice did not show any obvious differences gross morphology (A) body weight (A) or temperature (C) compared to age-matched litter mate control mice. **D**, survival curve shows that control mice (n = 57) survive to adulthood (30 days postnatal) while SCN1a^A1783V/+^ mice (n = 41) die prematurely starting at 9 days postnatal and reaching 100% lethality by 25 days (χ^2^ = 63.9, p < 0.0001). **E-F**, fluorescent *in situ* hybridization was performed to characterize expression of SCN1a transcript in inhibitory (VGAT+) and glutamatergic (VGLUT+) neurons in the RTN region in brainstem sections from control and SCN1a^A1783V/+^ mice. **E**, brainstem sections from control and SCN1a^A1783V/+^mice containing the RTN show SCN1a labeling (green puncta) of both VGAT+ and VGLUT neurons. **F**, summary data show SCN1a transcript expression (normalized to cell size) in VGAT+ and VGLUT+ RTN neurons from each genotype; channel transcript was reduced in VGAT+ cells from SCN1a^A1783V/+^ mice (0.43 ± 0.7 mRNA/area, n = 94 cells) compared to control (0.73 ± 0.9 mRNA/area, n = 82 cells) (p < 0.05), whereas VGLUT+ cells showed low channel transcript across both genotypes. These results were compared using a two-way ANOVA and Sidak multiple comparison test. *, p < 0.05; ***p < 0.001; ****p < 0.0001.

**Table 1.** Behavioral Assessment of Seizure activity.

| **Table 1: Behavioral Assessment of Seizure activity** |  |  |  |  |  |  |
| --- | --- | --- | --- | --- | --- | --- |
| Racine Score | 1 | 2 | 3 | 4 | 5 | Behavioral Arrest |
| Control | 5 | 0 | 0 | 0 | 0 | 0 |
| SCN1a<sup>A1783V/+</sup> | 3 | 3 | 6 | 2 | 3 | 5 |

To determine whether SCN1a^A1783V/+^ mice exhibit abnormal brain activity, we obtained electrocorticogram (ECoG) recordings from control and SCN1a^A1783V/+^ mice. We allowed mice 12 hours to recover after implanting them with the ECoG head stage. We continuously recorded ECoGs over a two-hour period, between the hours of 9:00 AM – 2:00 PM. Consistent with frequent polyspike activity observed in the ECoG recordings of DS patients (Bender et al., 2012), SCN1a^A1783V/+^ mice showed spontaneous high-amplitude spike-wave discharges (SWD) that occurred at a frequency of 0.37 ± 0.05/min (T_10_ = 3.009, p < 0.01) and had an average duration of 12.4 ± 5.5 s (Figs. 2A-B; T_10_ = 2.268, p < 0.05). Conversely, control animals showed SWD events less frequently (0.13 ± 0.1/min) and with shorter durations (6.7 ± 2.2s) compared to SCN1a^A1783V/+^ mice (Figure 2A; T_9_ = 2.4, p > 0.05). Power spectral analysis of SCN1a^A1783V/+^ SWD events showed increases in both alpha and beta frequency (F 4, 840 = 5.605, p < 0.001). These results suggest SCN1a^A1783V/+^ mice have frequent, spontaneous seizures and die prematurely (Figure 1D). Because this SUDEP-like phenotype is virtually identical to other DS models, including global and inhibitory neuron-specific SCN1a haplo-insufficient models (Yu et al. 2006, Kalume et al. 2013, Kim et al. 2018), we consider the SCN1a^A1783V/+^ mouse model to be useful for dissecting the mechanisms that underlie respiratory failure in DS.

**Figure 2.**
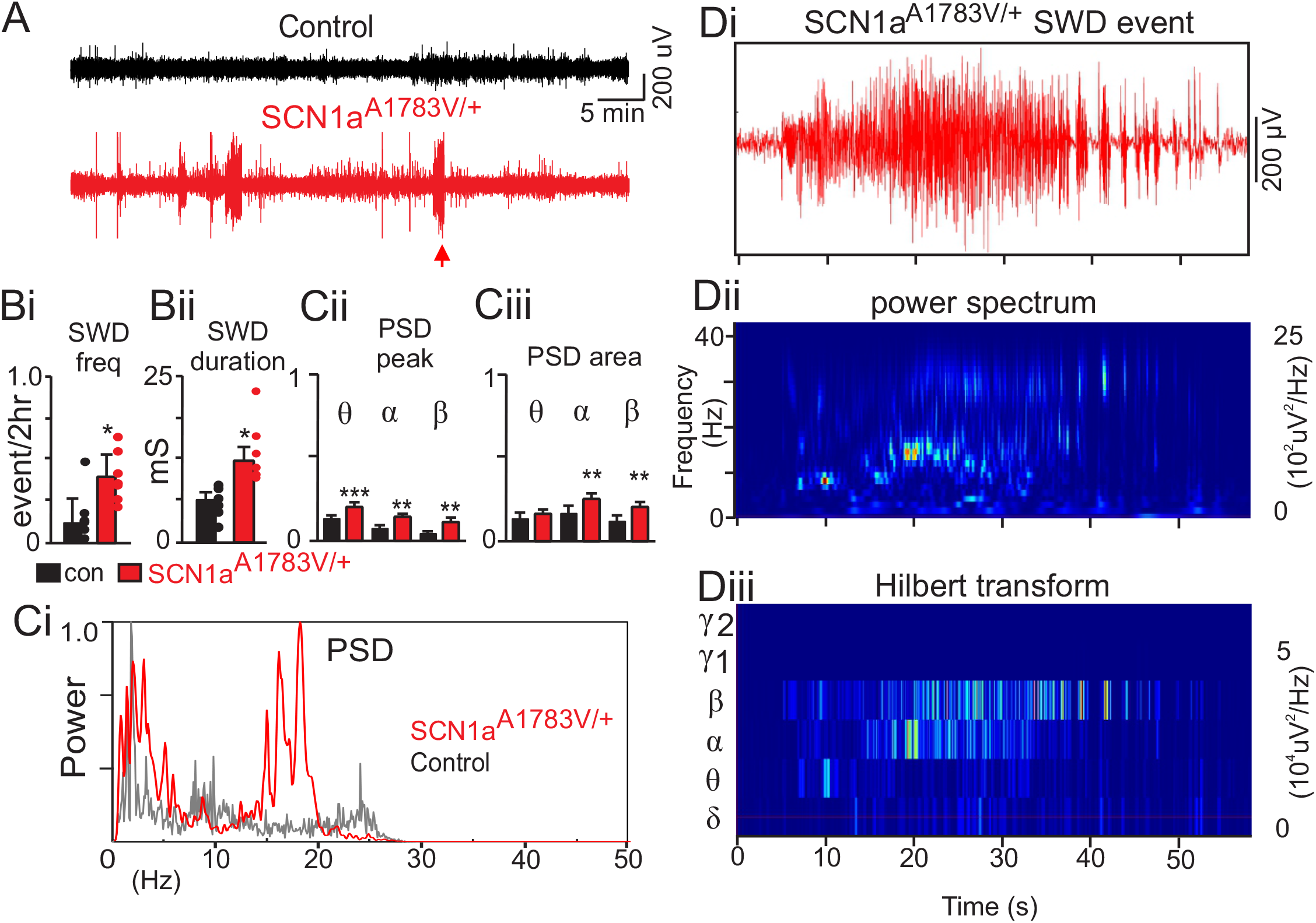
SCN1a^A1783V/+^ exhibit frequent spontaneous seizures. **A**, traces of raw EcoG activity show that SCN1a ^A1783V/+^ mice exhibit frequent spontaneous burst of high amplitude spike-wave discharges (SWD). The arrow identifies a typical seizure-like SWD event that was analyzed further by power spectral analysis in panel D. **B**, summary data show that seizure-like SWD events occurred more frequently (control 0.13 ± 0.1 events/2 hr, n = 6; SCN1a ^A1783V/+^ 0.37 ± 0.05 events/2 hr, n = 6; T 10=3.009, p < 0.01) and lasted for a longer duration (control 6.7 ± 2.2 ms, n = 6; SCN1a ^A1783V/+^ 12.4 ± 5.5 ms, n = 6, T 10=2.268, p < 0.05) in SCN1a ^A1783V/+^ mice compared to control animals. **Ci**, representative power spectrum density (PSD) plots of SWD events show typical strong activity in the theta-, alpha and beta frequency range in SCN1a^A1783V/+^ but not control mice. **Cii-Ciii**, summary data (normalized to the maximum value at each event) show PSD peak (Cii) and PSD area under the curve (Ciii) of each frequency range for each genotype. Note that SWD events measured in SCN1a^A1783V/+^ mice show increased activity in the theta, alpha and beta range but no differences in the delta or gamma band compared to SWD events measured in control mice. **Di-iii**, SWD event recorded from a SCN1a^A1783V/+^ mouse (arrow in panel A) plotted on an expanded time scale (Di) and corresponding time frequency distribution (Dii) and deconstructed spectrum into its various frequency domains (Diii). These results were compared using a two-way ANOVA and the Sidak multiple comparison test. *, p < 0.05; **, p< 0.01; ***p < 0.001.

### SCN1a^A1783V/+^ mice hypoventilate under baseline conditions and have a reduced CO_2_/H^+^ ventilatory response

Recent evidence (Kim et al., 2018) showed that DS patients have post-ictal respiratory abnormalities, including hypoventilation, apnea and impaired CO_2_ chemoreception. These symptoms can last for several hours after seizure, which indicates that respiratory problems contribute to SUDEP in DS. Therefore, to determine whether SCN1a^A1783V^ mice exhibit respiratory problems, we used a whole-body plethysmography to measure baseline breathing and the ventilatory response to CO_2_ in 15-day-old control and SCN1a^A1783V^ mice. We found that compared to control mice, SCN1a^A1783V^ mice showed a diminished respiratory output under room air conditions. Specifically, SCN1a^A1783V^ exhibited suppressed frequency (246 ± 15.3 bpm for controls compared to 199.5 ± 8.4 bpm for SCN1a^A1783V/+^; T_25_ = 2.665; p < 0.05); tidal volume (13.8 ± 2.0 µl/g for controls compared to 7.3 ± 1.7 µl/g for SCN1a^A1783V/+^, T_37_ = 2.351, p < 0.05); and minute ventilation (3.4 ± 0.5 µl/min/g for controls compared to 1.8 ± 0.4 µl/min/g for SCN1a^A1783V/+^; T_37_ = 2.173, p < 0.05; see Figures 3A-D for all). Although both genotypes exhibited apneic events at similar frequencies (0.23 ± 0.1/min for control and 0.11 ± 0.04/min for SCN1a^A1783V/+;^; p = 0.6), the duration of these events were longer in SCN1a^A1783V/+^ mice (Figure 3E; 1,104 ± 58.6 ms for controls versus 1,350 ± 99. 2 ms SCN1a^A1783V/+;^; T_51_ = 2.135; p < 0.05). We also found that SCN1a^A1783V/+^ mice had a diminished capacity to increase respiratory frequency in response to graded increases in CO_2_ (Figure 3F). Specifically, respiratory frequency in 7% CO_2_ (balance O_2_) was higher in controls (363.1 ± 7.7 bpm; N = 22) versus SCN1a^A1783V/+^ (300.7 ± 17.4 bpm; N = 17; F _1,37_ = 5.69, p < 0.05). Although tidal volume responses to CO_2_/H^+^ are similar between genotypes (p = 0.47), total respiratory output, as measured by minute ventilation— the product of respiratory frequency and tidal volume—was diminished in SCN1a^A1783V/+^ mice compared to controls (F3,111 = 3.167, p < 0.05; Figures 3G-H). Specifically, increasing inspired CO_2_ from 0% to 3% increased minute ventilation in control mice by 3.3 ± 0.5 µl/min/g (p < 0.0001). These same conditions, however, led to a much smaller and non-significant increase in minute ventilation among SCN1a ^A1783V/+^ mice (increase of 1.5 ± 0.5 µl/min/g; p = 0.07). These results show that SCN1a^A1783V/+^ mice exhibit a respiratory phenotype similar to that observed in DS patients, and further supports the possibility that respiratory problems may contribute to mortality in this DS model.

**Figure 3.**
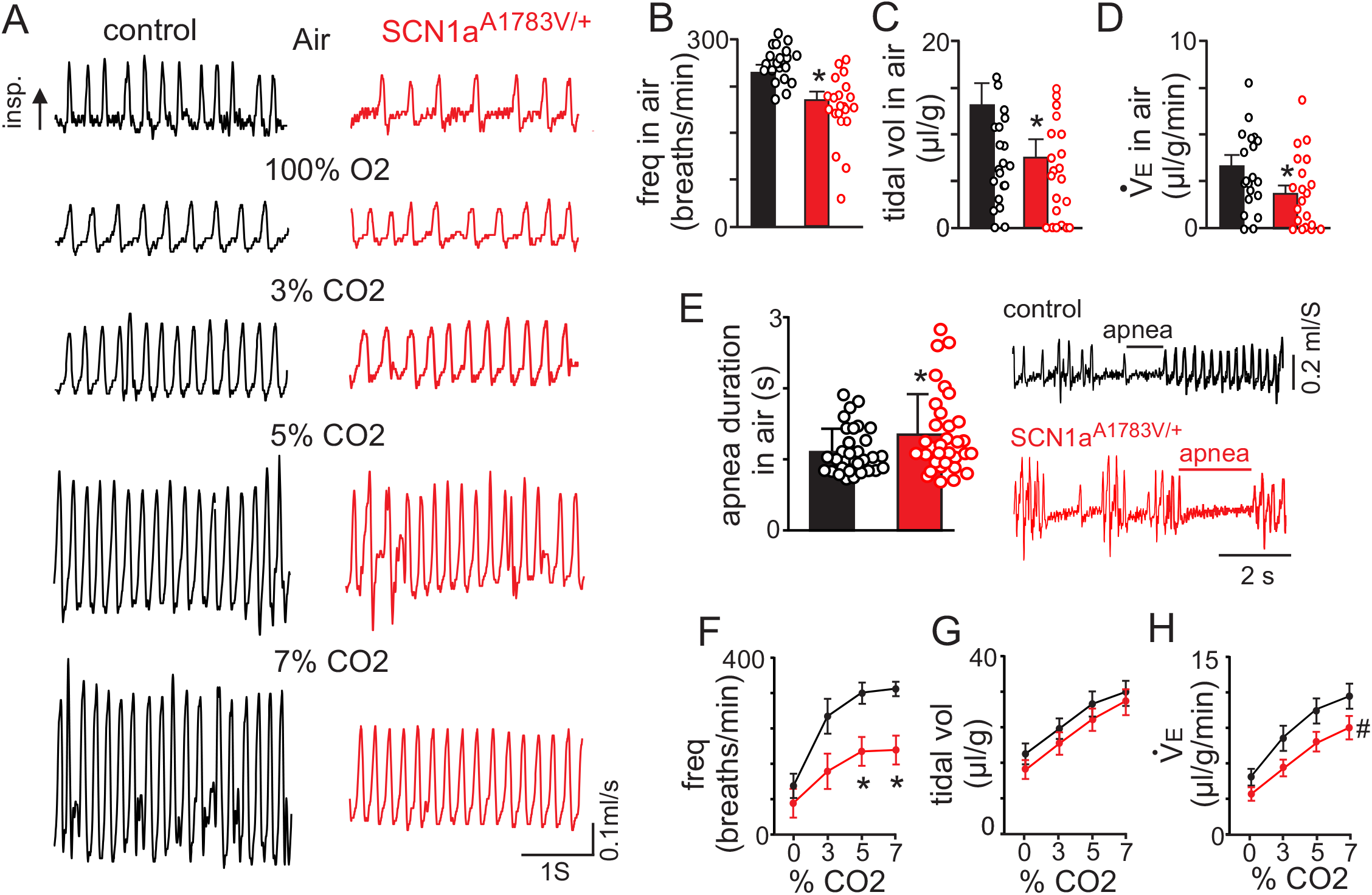
SCN1a^A1783V/+^ mice show reduced respiratory output under control conditions and during exposure to high CO_2_. **A**, traces of respiratory activity from a control and SCN1a^A1783V/+^ mouse during exposure to room air, 100% O2 and 3-7% CO2 (balance O_2_). **B-D**, summary data (n=22 control; n=17 SCN1a^A1783V/+^) show respiratory frequency (B), tidal volume (C) and minute ventilation (D) are reduced in SCN1A^A1783V/+^ mice compared to control under room air conditions. **E**, traces of respiratory activity (left) and summary data (right) show that under room air conditions both control and SCN1a^A1783V/+^ mice exhibit periods of apnea; the frequency of these events were similar between genotypes, however, they lasted for a longer duration in SCN1a^A1783V/+^ mice compared to control. **F-H**, summary data shows the respiratory frequency (F), tidal volume (G) and minute ventilation response of control and SCN1a^A1783V/+^ mice to graded increases in CO_2_ (balance O_2_). SCN1a^A1783V/+^ mice showed a blunted respiratory frequency to 5% and 7% CO_2_ which resulted in a diminished CO_2_/H^+^-dependent increase in minute ventilation. These results were compared using either unpaired t test (panels B-E) or two-way ANOVA followed by the Holm-Sidak multiple comparison test (panels F-H). *, difference between means p < 0.05, ^#^, different interaction factor, p < 0.05.

### Disinhibition and altered RTN chemoreception may underlie breathing problems in SCN1a^A1783V/+^ mice

The RTN is an important respiratory center and disrupting CO_2_/H^+^-sensitive cells in this region results in a respiratory phenotype similar to SCN1a^A1783V/+^ mice (Figure 3). Evidence also suggests that inhibitory neurons in the RTN region contribute to respiratory drive (Ott et al. 2011). We therefore sought to determine whether loss of SCN1a function in inhibitory neurons decreases inhibitory neuron activity and disinhibits excitatory, chemosensitive, neurons. To facilitate the identification of inhibitory neurons, we crossed VGATcre mice with the Rosa26^TdTomato^ reporter line. We crossed the resulting offspring with floxed-stop SCN1a^A1783V^ mice. VGAT+ cells that undergo this recombination express tdT and the A1783V variant (VGAT^tdT/+^:SCN1a^A1783V/+^). Consistent with other SCN1a knockout (Tai et al. 2014) or missense knockin (Ogiwara et al. 2007, Mashimo et al. 2010, Hedrich et al. 2014) DS models, the loss of SCN1a function in inhibitory neurons suppressed inhibitory neural activity. We performed whole-cell current clamp recordings of inhibitory neurons in the RTN region in VGAT^tdT/+^ slices, which yielded the following results: the inhibitory neurons of SCN1a^A1783V/+^ mice showed lower basal activity than those of controls (14.39 ± 1.5 Hz for controls vs. 9.902 ± 0.64 Hz for SCN1a^A1783V/+^; T_60_ = 2.97, p < 0.01; Figures 4A-B). Furthermore, SCN1a^A1783V/+^ inhibitory neurons fired fewer action potentials in response to depolarizing current steps (0-300 pA; ∆ 20 pA) from a holding potential of −80 mV. This activity deficit became more pronounced during large (200-300 pA) sustained (1,000 ms) current injections where inhibitory neurons expressing SCN1a^A1783V/+^ showed pronounced spike amplitude and frequency decrement (Figures 4A and D). That is, the number of spikes elicited by a +300 pA current step (1,000 ms) was 53.7 ± 11 for controls (N = 13) compared to 13.9 ± 6.4 for SCN1a^A1783V/+^ (N = 20; F15,465 = 9.536; p < 0.0001). We also found that inhibitory neurons from each genotype had similar input resistance (517.6 ± 82.2 M Ω for control vs. 519.2±38.9 M Ω for SCN1a^A1783V/+^; T_25_ = 0.02; p = 0.3; Figure 4C). These results suggest that expresson of SCN1a^A1783V/+^in brainstem inhibitory neurons reduces spontaneous activity and the neuron’s ability to respond to a wide range of excitatory inputs.

**Figure 4.**
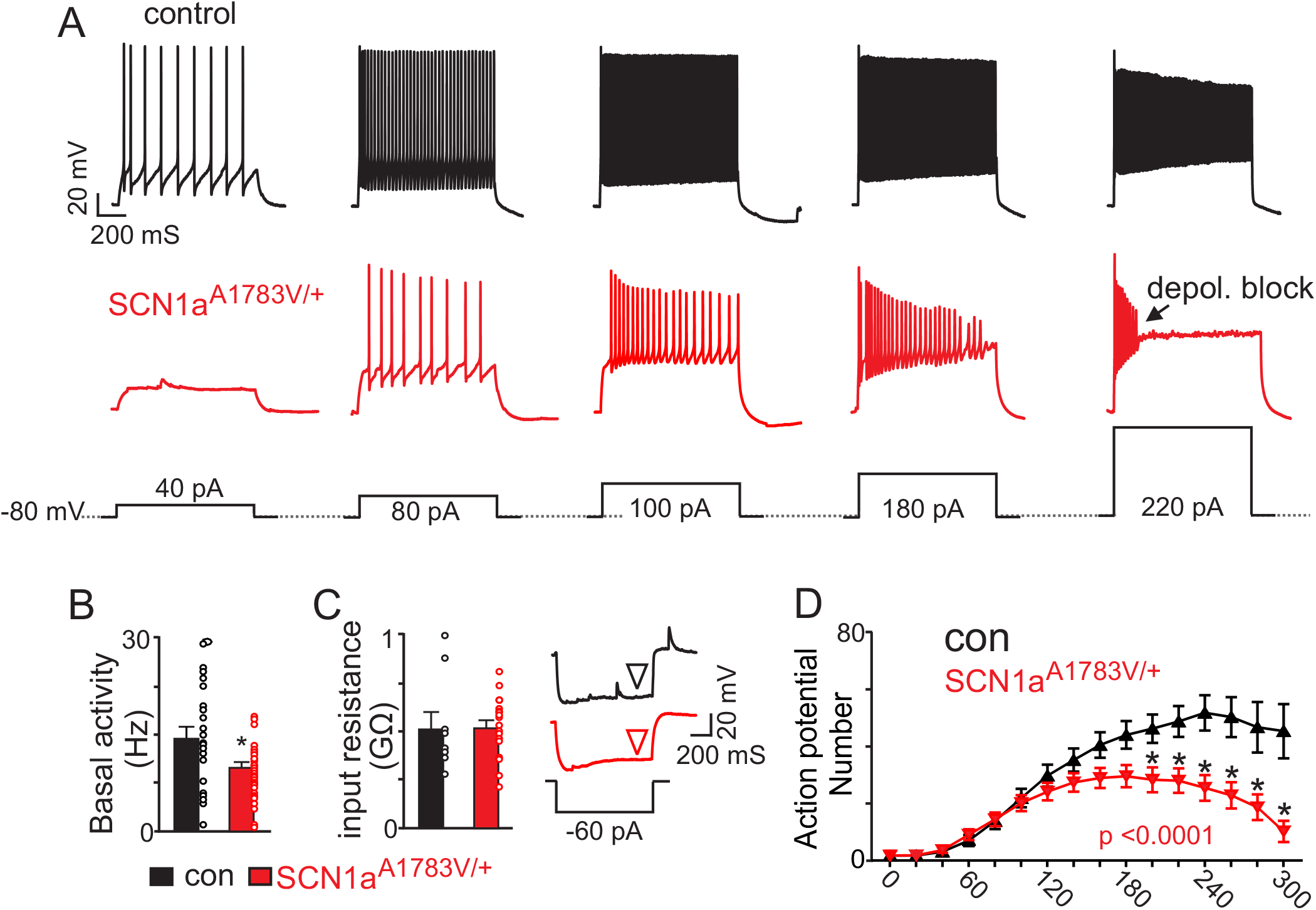
Brainstem inhibitory neurons in slices from SCN1a^A1783V/+^ show diminished basal activity and repetitive firing behavior during sustained depolarization. **A**, segments of membrane potential from inhibitory neurons in the RTN region in slices from control and SCN1a^A1783V/+^ mice during depolarizing current injections (40 to 220 pA; 1 s duration) from a membrane potential of –80 mV. **B**, summary data shows inhibitory neurons in slices from SCN1a^A1783V/+^ mice (n=36) are less active under resting conditions (0 pA holding current) compared to control cells (n = 26 cells). **C**, summary data and representative voltage responses to a −60 pA current injection show that inhibitory neurons from each genotype had similar input resistance. **D**, input-output relationship show that inhibitory neurons from SCN1a^A1783V/+^ mice generate fewer action potentials in response to moderate depolarizing current injections (1 s duration) and at more positive steps go into depolarizing block. Results were compared using t-test (B-C) and two-way ANOVA and Sidak multiple comparison test (D). *, p < 0.05; **, p<0.01; ***, p < 0.001.

The A1783V mutation is located in the S6 segment of domain 4 (Marini, Mei et al. 2007, Lossin 2009), a region thought to regulate voltage-dependent inactivation (Catterall WA 2000). Given that inhibitory neurons that express A1783V show reduced excitability, we hypothesized that the SCN1a A1783V variant results in loss of function by causing Nav1.1 channels to inactivate at more negative voltages. Consistent with this hypothesis, when examining spontaneous action potentials (as measured under resting conditions with a 0 pA holding current) in inhibitory neurons in slices of SCN1a^A1783V/+^ and control mice, the latter showed a higher amplitude (73.5 ± 1.9 mV) than SCN1a^A1783V/+^ (61.9 ± 2.6 mV; F _1,95_ = 9.931, p < 0.001). The maximum rate of depolarization was higher for controls (134.4 ± 5.0 mV/mS) compared to SCN1a^A1783V/+^ (89.4 ± 5.3 mV/mS; F_1,96_ = 35.2, p < 0.0001; see Figures 5A, D-F). Action potential threshold was also higher in inhibitory neurons in slices from SCN1a^A1783V/+^(−29.2 ± 0.9 mV) compared to controls (−32.4 ± 0.6 mV; F _1,95_ = 7.403, p < 0.05; Figures 5A, F).

**Figure 5.**
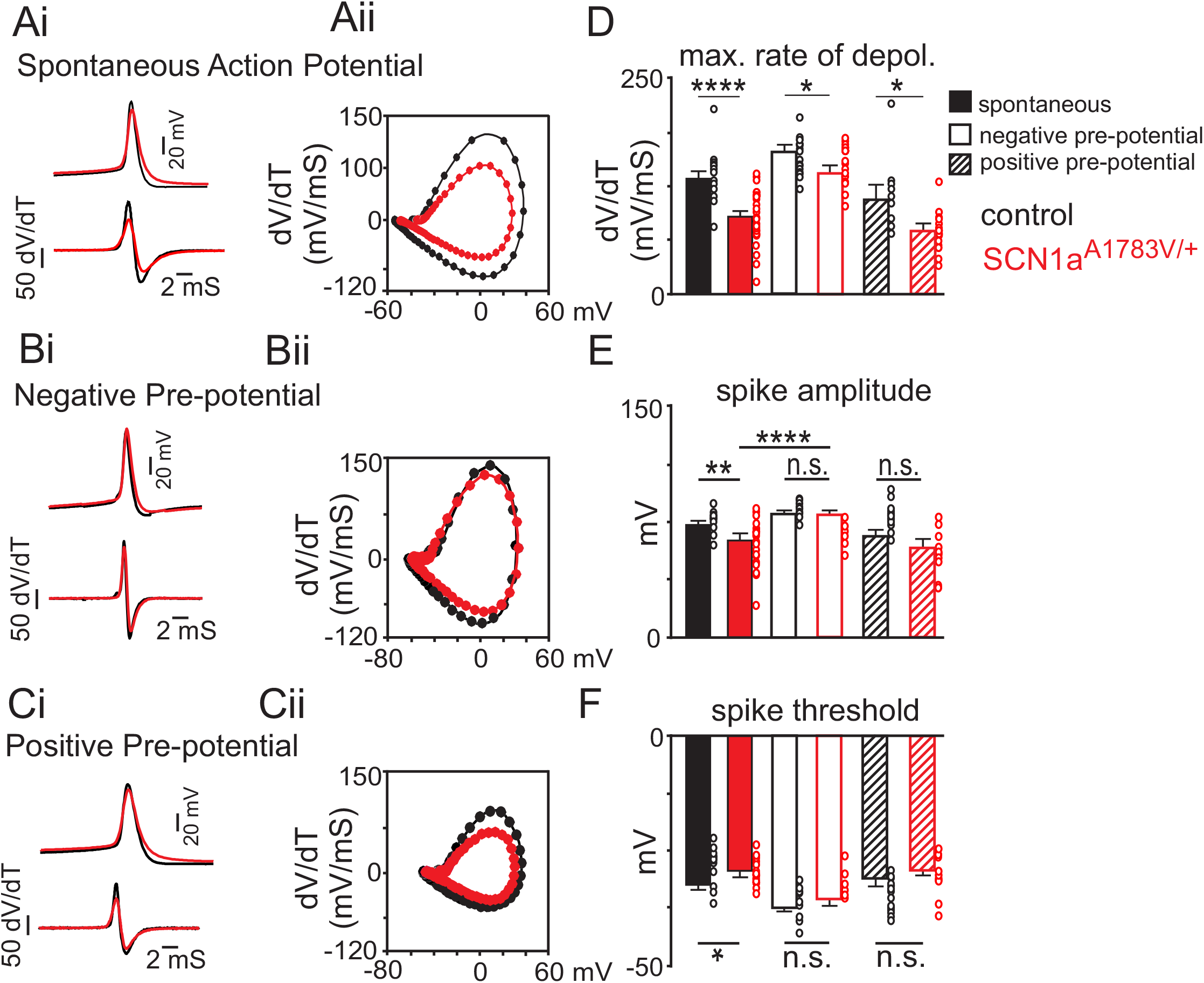
The SCN1a ^A1783V^ missense mutation appears to result in loss of channel function by increased voltage-dependent inactivation. **A**, average spontaneous action (control n= 24 spikes, SCN1a^A1783V/+^ n= 29 spikes (top) and first time derivative of average action potentials (bottom) recorded from inhibitory neurons in slices from control and SCN1a^A1783V/+^ mice (Ai) and corresponding phase plot (Aii) (dV/dt; Y-axis vs mV; X-axis) of the traces in panel Ai show that cells expressing SCN1a^A1783V/+^ depolarize slower compared to control cells. **B**, average first action potential following a hyperpolarizing pre-potential (−100 pA; 1 s) (control n= 13 spikes, SCN1a^A1783V/+^ n= 16 spikes) (top) and first time derivative of average action potentials (bottom) recorded from inhibitory neurons in slices from control and SCN1a^A1783V/+^ mice (Bi) and corresponding phase plot (Bii) of traces in panel Bi show that holding cells at a negative pre-potential to remove sodium channel inactivation improved the depolarization kinetics of subsequent spikes. **C**, average first action potential following a depolarizing pre-potential (+180 pA; 1 s) (control n= 9 spikes, SCN1a^A1783V/+^ n= 11 spikes (top) and first time derivative of average action potentials (bottom) recorded from inhibitory neurons in slices from control and SCN1a^A1783V/+^ mice (Ci) and corresponding phase plot (Cii) of traces in panel Ci show that holding cells at a depolarized pre-potential to increase sodium channel inactivation diminished genotype differences in action potential kinetics. **D-F**, summary data showing the maximum rate of depolarization (D), action potential amplitude (E) and action potential threshold (F) of spontaneous action potentials and first spikes following positive or negative pre-potentials recorded in slices from control and SCN1a^A1783V/+^ mice. Results were compared by two-way ANOVA and the Sidak multiple comparison test.. *, p < 0.05; **, p< 0.01; ***, p < 0.001; ****, p < 0.0001.

Next, we characterized the properties of the first action potential elicited after holding cells at potentials that either remove or enhance Nav1.1 channel inactivation. We found that differences in action potential waveform properties between genotypes were minimized when we held cells at a negative voltage to remove Na^+^ channel inactivation. For example, holding inhibitory neurons expressing SCN1a^A173V/+^ at a negative pre-potential by injecting a hyperpolarizing current (−100 pA; 1,000 ms) increased action potential amplitude (78.01 ± 2.0 mV SCN1a^A1783V/+^; F_1, 95_ = 9.931, p < 0.0001) to an amount similar to spikes from control cells (81.05 ± 1.2 mV control; p = 0.83) (Figures 5B, E). Under these conditions, the maximum rate of depolarization also increased 140.3 ± 5.441 mV/ms (F _1, 96_ = 35.21, p < 0.0001) (Figures 5B, D); this rate was similar to that measured in spontaneous spikes from control animals (p > 0.99) but slower than spikes from control cells following a negative pre-potential (166.6 ± 7.3 mV/mS, F _1,96_ = 35.21, p < 0.05). Holding inhibitory neurons expressing SCN1a^A1783V/+^ at a negative pre-potential also lowered the threshold for spike initiation (−35.68 ± 0.7 mV SCN1a^A1783V/+^; F_1, 95_ = 7.403, p<0.001) to a level similar to control cells (−37.2 ± 1.2 mV control; F_1, 95_ = 7.403, p = 0.06) (Figures 5B, F). We also found that delivering a +180 pA current for 1,000 ms to enhance Na^+^ channel inactivation in control cells resulted in similar action potential amplitude (F_1, 95_ = 9.931, p > 0.99), rate of depolarization (F_1, 96_ = 35.21, p = 0.58) and spike threshold (F_1, 95_ = 7.403, p = 0.97) as spikes measured in inhibitory neurons from SCN1a^A1783V/+^ slices under resting conditions (holding current = 0 pA) (Figures 5C, D-F). Together, these results are consistent with the possibility that the SCN1a A1783V mutation results in loss of function due to enhanced Nav1.1 inactivation.

Previous evidence suggests that inhibitory neurons in the RTN region regulate activity of chemosensitive neurons (Ott et al. 2011). Because expression of SCN1a^A1783V^ suppresses activity of inhibitory neurons in the RTN region (Figures 4 and 5), we predict that loss of inhibitory tone would enhance basal activity and CO_2_/H^+^ sensitivity of excitatory chemosensitive neurons. To test this, we characterized the firing activity of chemosensitive RTN neurons in slices from each genotype during exposure to CO_2_ levels ranging from 3 to 10%. We initially identified chemosensitive RTN neurons in each genotype by their firing response to CO_2_. We considered neurons that were spontaneously active in 5% CO_2_ and responded to 10% CO_2_ with at least a 1.0 Hz increase in firing rate to be chemosensitive. Chemosensitive RTN neurons also have been shown to express the transcription factor Phox2b; therefore, at the end of each experiment we filled all recorded cells with Lucifer yellow for later immunohistochemical confirmation of Phox2b expression. Chemosensitive RTN neurons in slices from control mice had an average basal activity of 1.3 ± 0.4 Hz under control conditions (5% CO_2_; pH 7.3). These cells were strongly inhibited by decreasing CO_2_ to 3% (pHo = 7.6) (1.02 ± 0.3 Hz) and showed a linear firing increase in response to 7% (pHo = 7.2) (2.4 ± 0.5 Hz) and 10% CO 2 (pHo = 7.0) (2.8 ± 0.4 Hz) (Figures 6A, B-C). This CO 2 response profile is consistent with type I chemoreceptors (pH_50_= 7.3), which were described previously in a Phox2b mouse reporter line (Lazarenko et al. 2009). Consistent with our hypothesis, chemosensitive RTN neurons in slices from SCN1a^A1783V/+^ mice were more active under control conditions (5% CO_2_) (2.4 ± 0.35 Hz)(Figure 6C) (T_21_=2.223, p < 0.05) and showed an enhanced firing response to high CO_2_/H^+^ (Figure 6D) (slope: 0.3 ± 0.01 control vs. 0.37 ± 0.01, F1,4 =8.04, p < 0.05). These results show that loss of SCN1a function in inhibitory neurons disrupts activity of RTN chemoreceptors.

**Figure 6.**
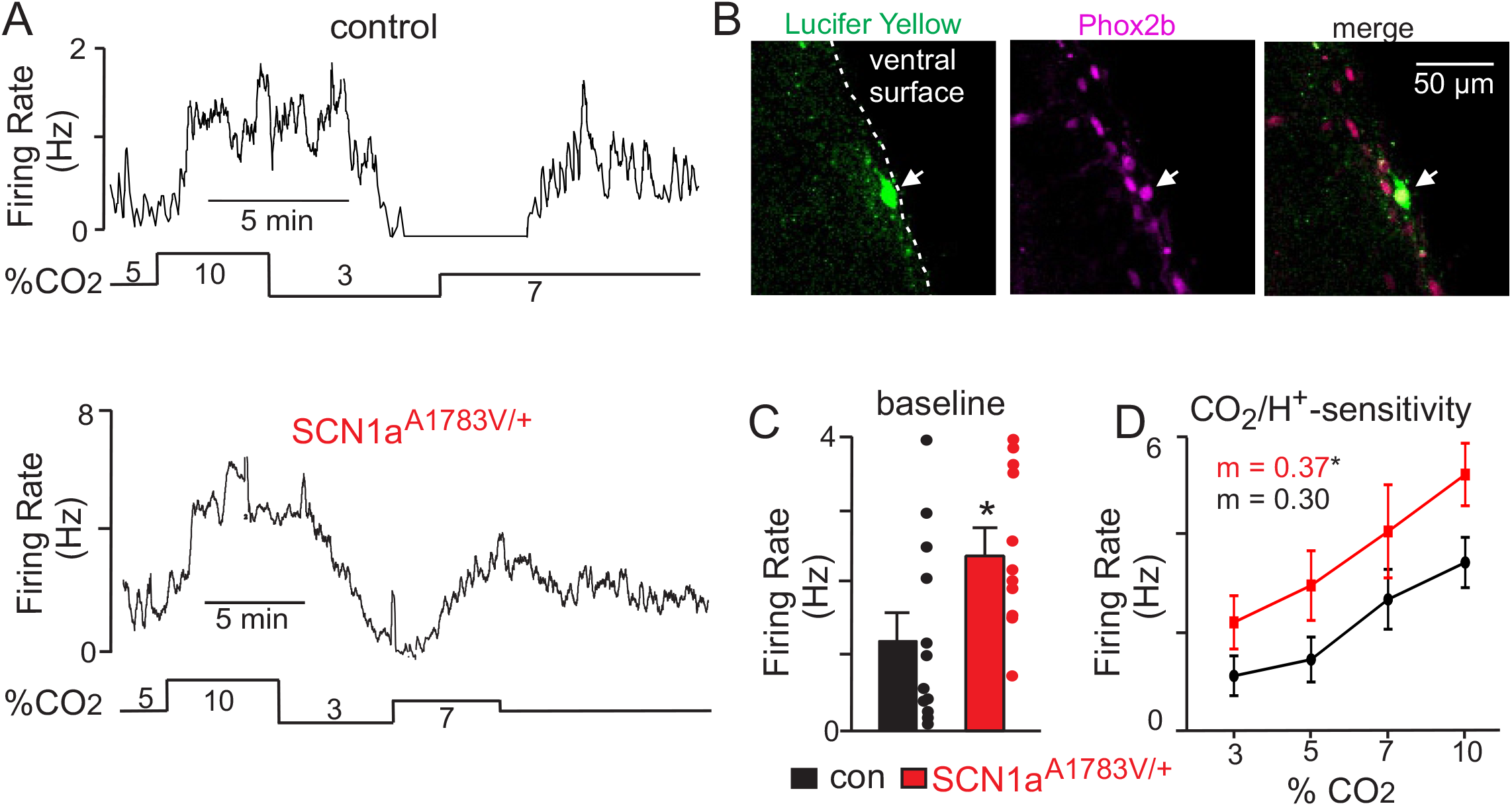
Chemosensitive RTN neurons in slices from SCN1a^A1783V/+^ mice are hyper-excitable. **A**, firing rate traces from chemosensitive neurons in slices from control (top) and SCN1a^A1783V/+^ mice (bottom) show that neurons from both genotypes respond to changes in CO_2_/H^+^; RTN neurons are spontaneously active under control conditions (5% CO_2_; pHo 7.3) and respond to 7% CO_2_ (pHo 7.2) and 10% CO_2_ (pHo 7.0) with a linear increase in activity, whereas exposure to 3% CO_2_ (pHo 7.6) decreases neural activity. However, basal activity and CO_2_/H^+-^ dependent output of RTN chemoreceptors from SCN1a^A1783V/+^ tissue is enhanced compared to control cells. **B**, double-immunolabeling shows that a Lucifer Yellow-filled CO_2_/H^+^-sensitive RTN neuron (green) is immunoreactive for phox2b (magenta), the merged image is shown to the right. We confirmed that all CO_2_/H^+^-sensitive neurons (control n= 12; SCN1a^A1783V/+^ n = 11) included in this study were phox2b-positive. **C-D**, summary data shows that RTN chemoreceptors in slices from SCN1a^A1783V/+^ mice have higher basal activity (C) and enhanced CO_2_/H^+^-dependent output between 3-10% CO_2_ (D). Results were compared by t-test (C) or ANCOVA test (D). *, p < 0.05.

## DISCUSSION

Epilepsy patients have a 40-fold higher mortality rate than the general population (Dlouhy et al. 2016). The most common cause of death for this patient population is SUDEP, a leading cause of which is respiratory failure (Surges et al. 2009, Ryvlin et al. 2013, Kennedy and Seyal 2015, Dlouhy et al. 2016). However, the mechanisms underlying respiratory dysfunction in epilepsy and SUDEP are largely unknown. This is particularly true in the context of DS, where patients have an exceedingly high mortality rate and commonly exhibit life-threatening respiratory problems (Kim et al. 2018), yet nothing is known regarding how loss of SCN1a function impacts brainstem respiratory centers. The results presented here address this knowledge gap by showing that expression of the recurrent DS-associated SCN1a variant A1783V in inhibitory neurons results in seizures and pre-mature death (Figures 1D, 2). Moreover, this mouse model presents with a respiratory phenotype strikingly similar to that exhibited by DS patients. Perhaps not surprising, we found that loss of SCN1a function in inhibitory neurons in the RTN diminished activity in a cell-autonomous manner but, importantly, also enhanced baseline activity and CO_2_/H^+^ sensitivity of excitatory chemosensitive neurons. These results suggest that disruption of SCN1a in inhibitory neurons can impact brainstem respiratory centers and contribute to pathological features of DS including disordered breathing associated with SUDEP

By ~ 2 weeks of age, SCN1a^A1783V^ mice exhibited a respiratory phenotype similar to DS patients including hypoventilation, increased apneas and diminished ventilatory response to CO_2_/H^+^. These breathing problems occurred in conjunction with a marked increase in mortality, thus correlatively supporting the possibility that respiratory failure contributes to premature death in DS. Although the mechanisms contributing to respiratory dysfunction in DS are unknown, previous work showed that loss of SCN1a from inhibitory neurons in the forebrain, but not the brainstem where respiratory control centers are located, resulted in premature death (Cheah et al. 2012). These results are consistent with the possibly that respiratory dysfunction is a secondary consequence of cortical seizure activity propagating to and disrupting brainstem function.

There are numerous direct and indirect projections from the cortex to brainstem respiratory centers (Shea 1996) that may serve as the anatomical substrate for seizure-induced respiratory dysfunction. For example, recent work in humans showed that apnea and arterial oxygen desaturation occurred when cortical seizure activity spread to the amygdala (Dlouhy et al. 2015) and presumably activated descending inhibitory projections to brainstem respiratory centers. However, SUDEP can also occur in epilepsy patients in the absence of an overt seizure or outside the peri-ictal period (Lhatoo and Shorvon 1998), suggesting that factors other than acute seizure predispose individuals to SUDEP. For example, it is possible that repeated bombardment of brainstem respiratory centers by frequent cortical seizure events alters cellular or neural network function, leading to progressive respiratory disruption and increased SUDEP propensity. Consistent with this possibility, patients with temporal lobe epilepsy (a common type of focal epilepsy) show widespread alterations in neural network activity including at the level of the brainstem (Englot et al. 2018). However, it remains unclear whether elements of respiratory control are compromised by repeated seizure activity in a similar manner.

Our results show that SCN1a transcript is highly expressed by brainstem inhibitory neurons and to a lesser extent by glutamatergic neurons (Figures 1E-F); therefore, loss of SCN1a function is likely to directly impact brainstem inhibitory neurons independent of descending seizure activity. Consistent with this possibility and analogous to cortical inhibitory neurons in SCN1a^-/+^ knockout models (Cheah et al. 2012) and induced pluripotent stem cells derived from DS patients with an SCN1a truncation mutation (Higurashi et al. 2013), we found that inhibitory neurons in the RTN region expressed the SCN1a A1783V variant produced fewer action potentials in response to sustained depolarizing current injection and were more prone to depolarization block compared to inhibitory neurons from control mice (Figures 4A-D). These results suggest that loss of SCN1a might suppress inhibitory tone in brainstem respiratory centers including the RTN where inhibitory neurons appear to interact with and regulate the activity of excitatory chemosensitive neurons (Ott et al. 2011).

We confirmed this possibility at the cellular level by showing that baseline activity and CO_2_/H^+-^dependent output of RTN chemoreceptors in slices from SCN1a^A1783V/+^ mice were enhanced compared to RTN chemoreceptors in slices from control mice, thus demonstrating that RTN chemoreceptor function is altered in this DS model. However, because chemosensitive RTN neurons provide an excitatory drive to breathe (Guyenet et al. 2016), we would predict that disinhibition of these neurons would potentiate rather than suppress respiratory function as observed in DS patients (Kim et al. 2018) and SCN1a^A1783V/+^ mice (Figure 3). Therefore, it is likely that other respiratory elements also contribute to the observed respiratory phenotype. In particular, inhibitory signaling within the pre-Böt zinger complex (pre-BötC) – a brainstem region downstream of the RTN that regulates inspiratory rhythm – is required for rapid breathing; Studies have reported that disrupting inhibitory neuromodulation within this region prolongs the post-burst refractory period exhibited by pre-BötC inspiratory neurons, resulting in reduced respiratory frequency (Baertsch et al. 2018). Therefore, it is reasonable to speculate that global loss of SCN1a function could overactivate pre-BötC inspiratory neurons and increase the inspiratory burst refractory period, thus slowing respiratory frequency. Furthermore, as chemosensitive RTN neurons send excitatory glutamatergic projections directly to inspiratory pre-BötC neurons, disinhibition at the level of the RTN would likely further compromise inspiratory output by the pre-BötC.

Despite the prevalence of SCN1a missense mutations in DS (Parihar and Ganesh 2013), few studies have characterized the pathophysiology associated with specific mutant alleles. This is particularly important for the development of patient-directed therapies because the aberrant products of SCN1a missense mutations are potentially expressed, thus representing a novel therapeutic target that is absent from haploinsufficient models of DS. Here, we show that expression of a DS-associated missense mutation (A1783V) in inhibitory neurons results in seizures and premature death on a similar time scale as haploinsufficient DS models (Caterall WA 2012). Inhibitory neurons from SCN1a^A1783V/+^ mice showed a modest reduction in channel transcript that may contribute to loss of inhibitory tone; however, the repetitive firing characteristics of SCN1a^A1783V/+^-expressing neurons was more consistent with loss of function due to increased Nav1.1 channel inactivation. For example, genotype differences in the action potential amplitude and rate of depolarization were diminished under experimental conditions designed to remove Na^+^ channel inactivation. Therefore, an effective treatment for SCN1a^A1783V/+^-associated pathology could be to selectively potentiate Nav1.1 channel activity by slowing voltage-dependent inactivation. This approach, using a spider venom called heteroscodratoxin-1 (Hm1a), has been show to decrease seizures and mortality in an SCN1a haploinsufficient model of DS, albeit at a concentration well above the Hm1a EC_50_ of Nav1.1 channels (Richards et al., 2018). If the potency of Hm1a depends on channel availability, then we predict that Hm1a would rescue the function of inhibitory neurons expressing A1783V mutant channels more effectively than neurons expressing only one functional SCN1a allele. However, limitations associated with Hm1a, including poor blood-brain barrier permeability and a short half-life, preclude testing this possibility in a clinically relevant manner at this time.

In sum, our results show that expression of SCN1a^A1783V^ in inhibitory neurons mirror clinical features of DS including spontaneous seizures and respiratory dysfunction. At the cellular level, brainstem inhibitory neurons in the RTN of slices from SCN1a^A1783V^ are less excitable whereas glutamatergic chemosensitive neurons are more excitable. Thus, our findings indicate that RTN chemoreceptors are a potential substrate for respiratory dysfunction in DS.

## ACKNOWLEDGEMENTS

We thank Drs. Ana Mingorance (Chief Development Officer of the Loulou Foundation) and Anastasios Tzingounis (Dept. Physiology and Neurobiology, Univ. Connecticut) for their constructive suggestions regarding this project. This work was supported by funds from the National Institutes of Health Grants HL104101 (DKM), HL137094 (DKM) and NS104999 (JLL). Additional funds were also provided by the Dravet Foundation Grant AG180243 (DKM) and American Epilepsy Society (F-SK).

## AUTHOR CONTRIBUTIONS

F-SK: experimental design; collection and analysis of data; revising the manuscript; final approval of the manuscript.

JJL: experimental design; revising the manuscript; final approval of the manuscript.

XC: data analysis; revising the manuscript, final approval of the manuscript.

DKM: experimental design; data analysis; drafting the manuscript; revising the manuscript, final approval of the manuscript.

## DECLARATION OF INTERESTS

The authors declare no competing interests

## METHODS

### SCN1a ^A1783V/+^ mice

All animal use was in accordance with guidelines approved by the University of Connecticut Institutional Animal Care and Use Committee. SCN1a ^A1783V /+^ mice were generated by crossing offspring of VGAT-ires-cre (JAX no. 016962) and tdTomato mice (JAX no. 007914) with mice carrying cre-driven heterozygous SCN1a A1783V mutation (Jax no. 026133) to introduce the SCN1a variant A1783V conditionally in inhibitory neurons. Experimental animals express both the reporter and the SCN1a^A1783V/+^ mutation (SCN1a ^A1783V/+^ mice), whereas litter mate control animals were those that express the tdTomato reporter under the VGAT promotor. Aged matched mice from both genotypes and sexes were used for all experiments included in this study.

### Fluorescent in situ hybridization (FISH)

To prepare fresh frozen slice, postnatal week 2 mice of both genotypes were anesthetized with isoflurane, decapitated, and brainstem tissues were rapidly frozen with dry ice and embedded with OCT compound. Brainstem slices (14um thick) containing the retrotrapezoid nucleus (RTN) were crysectioned and collected onto SuperFrost Plus microscope slides. Slices were fixed with 4% paraformaldehyde and dehydrated with 50%,70% and 100% ethanol. FISH were processed with the instruction of RNAscope^®^ Multiplex Fluores cent Assay (ACD, 320850), the probes used in our study were designed and validated by ACD (Table 2).

**Table 2.**
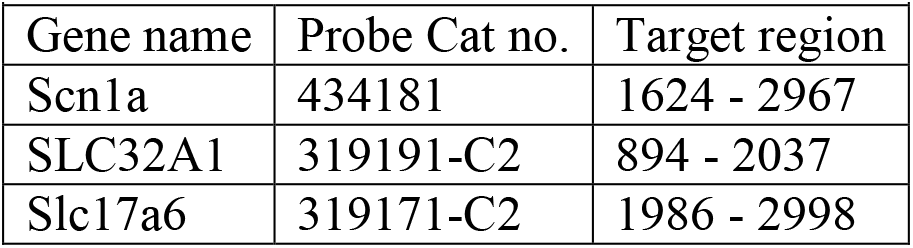
Probes used for FISH.

### Unrestrained whole-body plethysmography

Respiratory activity was measured using a whole-body plethysmograph system (Data Scientific International; DSI), utilizing animal chamber (600 mL volume) maintained at room temperature and ventilated with air (1.3 L/min) using a small animal bias flow generator. Fifteen day old mice (~7 g) were individually placed into a chamber and allowed 2 hour to acclimate prior to the start of an experiment. Respiratory activity was recorded using Ponemah 5.20 software (DSI) for a period of 15 minutes in room air followed by exposure to graded increases in CO_2_ from 0% to 7% CO_2_ (balance O_2_). Body temperature was measured before and after each experiment and although body temperature tended to drop ~ 1 ◦C by the end of an experiment, there were no genotype difference in the degree of cooling (p = 0.37). Parameters of interests include respiratory frequency (F_R_, breaths per minute), tidal volume (V_T_, measured in mL; normalized to body weight and corrected to account for chamber and animal temperature, humidity, and atmospheric pressure), and minute ventilation (V_E_, mL/min/g). A 20 second period of relative quiescence after 2 minutes of exposure to each condition was selected for analysis. Spontaneous apneic events, conservatively defined as 3 or more missed breaths not preceded by a sigh or augmented breath, were analyzed off-line. All experiments were performed between 9 a.m. and 6 p.m. to minimize potential circadian effects.

### Acute slice preparation and in vitro electrophysiology

Slices containing the RTN were prepared as previously described (Mulkey et al., 2007). In short, rats were anesthetized by administration of ketamine (375 mg/kg, I.P.) and xylazine (25 mg/kg; I.P.) and rapidly decapitated; brainstems were removed and transverse brain stem slices (300 µm) were cut using a microslicer (DSK 1500E; Dosaka) in ice-cold substituted Ringer solution containing the following (in mM): 260 sucrose, 3 KCl, 5 MgCl_2_, 1 CaCl_2_, 1.25 NaH_2_PO_4_, 26 NaHCO_3_, 10 glucose, and 1 kynurenic acid. Slices were incubated for 30 min at 37°C and subsequently at room temperature in a normal Ringer’s solution containing (in mM): 130 NaCl, 3 KCl, 2 MgCl_2_, 2 CaCl_2_, 1.25 NaH_2_PO_4_, 26 NaHCO_3_, and 10 glucose. Both substituted and normal Ringer’s solutions were bubbled with 95% O_2_ and 5% CO_2_ (pH=7.30).

Individual slices containing the RTN were transferred to a recording chamber mounted on a fixed-stage microscope (Olympus BX5.1WI) and perfused continuously (~2 ml/min) with a bath solution containing (in mM): 140 NaCl, 3 KCl, 2 MgCl_2_, 2 CaCl_2_, 10 HEPES, 10 glucose (equilibrated with 5% CO_2_; pH=7.3). All recordings were made with an Axopatch 200B patch-clamp amplifier, digitized with a Digidata 1322A A/D converter, and recorded using pCLAMP 10.0 software (Molecular Devices, Sunnyvale, CA). Recordings were obtained at room temperature (~22° C) with patch electrodes pulled from borosilicate glass capillaries (Harvard Apparatus, Molliston, MA) on a two-stage puller (P-97; Sutter Instrument, Novato, CA) to a DC resistance of 5–7 MΩ when filled with a pipette solution containing the following (in mM): 125 K-gluconate, 10 HEPES, 4 Mg-ATP, 3 Na-GTP, 1 EGTA, 10 Na-phosphocreatine (uM), 0.2% Lucifer yellow (pH 7.30). Electrode tips were coated with Sylgard 184 (Dow Corning, Midland, MI).

The firing response of chemosensitive RTN neurons to CO_2_ (3-10% CO_2_) was assessed in the cell-attached voltage-clamp configuration (seal resistance > 1 GΩ) with holding potential matched to the resting membrane potential (V_hold_ = −60 mV) and with no current generated by the amplifier (*I*_amp_ = 0 pA). Firing rate histograms were generated by integrating action potential discharge in 10 to 20-second bins using Spike 5.0 software (Cambridge Electronic Design, CED, Cambridge, U.K.). We confirmed that all chemosensitive RTN neurons included in this study were immunoreactive for the transcription factor Phox2b.

To characterize action potential properties and repetitive firing behavior of inhibitory neurons, we made whole-cell current-clamp recordings from fluorescently labeled neurons located in the region of the RTN in slices from VGAT:TdTomato mice. Repetitive firing responses to 1 second depolarizing current steps from 0 to 300 pA (∆ 20 pA increments) were characterized from an initial holding potential of −80 mV. Action potential amplitude, threshold (dV/dT > 10mV/mS) and the maximum rate of depolarization obtained from the peak of the first time derivative of the action potential were characterized for spontaneous spikes measured under resting conditions (holding current = 0 pA) and for the first spike elicited after delivering a positive (+200 pA) or negative (−100 pA) 1 second current step. All whole-cell recordings had an access resistance (Ra) < 20 MΩ, recordings were discarded if Ra varied >10% during an experiment. A liquid junction potential of −14 mV was accounted for during each experiment.

### Immunofluorescence staining

Slices were fixed in 4% PFA over night after recording, and blocked with 5% normal horse serum in 1X PBS with 2.5% triton for 1 hr. Slices were incubated in goat anti-phox2b antibody (R&D, AF4940) and rabbit anti-lucifer yellow antibodies (Invitrogen, A-5750) mixed in blocking solution under 4℃ overnight. After washing the primary antibody a secondary antibody was applied for 2 hrs followed by an additional wash and mounting with ProLong^®^ Gold Antifade Reagent (Invitrogen, P36934). Slices were imaged using a Leica SP8 confocal microscope (40x/1.3 HC oil objective) to identify cells that co-express Alex Fluro 647 for phox2b and Alex Fluro 488 for lucifer yellow.

### Electrocorticography recording

Subdural EcoG electrodes were implanted in 15 day old control and SCN1a^A1783V/+^ mouse pups. To minimize damage we used stainless steel wire electrodes (diameter = 0.003 in) (A-M system, 790900) inserted just under the skull for a length of 2 mm into each hemisphere near the fontal cortex. A reference wire electrode was placed in the posterior cortex. Each electrode was connected to a Mill-MAX miniature socket (digikey, ED11265-ND) and secured to the skull with super glue. Differential voltage signals were amplified 1000× with a DAM-50 differential amplifier (1 Hz low filter, 10 kHz high filter), digitized at 5 kHz and recorded using Sirenia Software (Pinnacle technology).

Mice were allowed to 12 hours to recover from surgery before recording EcoG activity for a period of 2 hours. We also video recorded all experiments to correlate animal behavior with EcoG recordings. Only spike wave discharge (SWD) activity that occurred in conjunction with observable seizure behavior was included in the analysis. Any data including movement artifacts were excluded from analysis. The same criteria for seizure events were used for both mutant mice and control group. The full duration of each seizure event was segmented and then down sampled from 600 Hz to 100 Hz to focus on the frequency range of interest (0-50 Hz) prior to performing the power spectral analysis in Matlab (MathWorks). Frequency ranges of EcoG signals are defined as follows: delta, δ (1–5 Hz), theta, Ө (6–8 Hz), alpha, α (9–16 Hz), beta, β (17–36 Hz), and gamma, γ (37–50 Hz). Frequency analysis results were normalized to the maximum frequency amplitude at each event. For each frequency range, maximal amplitude and area under each frequency range were calculated to report the spectral power. To show the time-varying frequency distribution, time frequency analysis using Hilbert and Morlet transformations were also performed using Brainstorm 3.0 (Tadel et al. 2011).

### Seizure behavior Scoring

We video monitored mice for 1 hour after placing them individually in a cage and giving them access to food and water ad lib. Seizure behavior during this time was evaluated using the Racine scoring system as follows: score 1, mouth and facial movements; score 2, head nodding; score 3, forelimb clonus; score 4, rearing with limb clonus; score 5, full body clonus, rearing and falling.

### Statistical Analysis

Data are reported as mean ± SE. All statistical analysis was performed using Prism 7 (GraphPad Software, Inc., La Jolla, CA). Data were normally distributed (Shapiro-Wilk normality test) and comparisons were made using t-test, Chi Square test, one-way or two-way ANOVA followed by multiple comparison tests as appropriate. Relevant values used for statistical analysis are included in the results section.

